# Brief visual oddball events evoke CSF flow in the spinal canal

**DOI:** 10.64898/2026.11.05.764672

**Authors:** Elizabeth Riley, Henning Voss, Eve De Rosa, Adam Anderson

**Affiliations:** Cornell Joan Klein Jacobs Center for Precision Nutrition and Health; Cornell MRI Facility, Cornell University; Department of Psychology, Cornell University

**Author notes:** Corresponding author: Elizabeth Riley.

**Keywords:** cerebrospinal fluid, CSF, fMRI, aging, multi-echo, oddball

## Abstract

The measurement of cerebrospinal fluid (CSF) flow with fMRI, made possible by exploiting edge artifacts of spin saturation when the bottom acquisition slice lies in a CSF reservoir, allows researchers to investigate an aspect of brain health that has been increasingly linked to cognitive aging and neurodegenerative disease via its role in glymphatic clearance. The initial demonstration of this technique used the fourth ventricle, but recent reports show it can also be done when the bottom acquisition slice lies in the spinal canal in the foramen magnum. We present a method to automatically identify the spinal canal without manual tracing, and measure CSF flow with high confidence using multi-echo acquisition to separate out BOLD-like and non-BOLD-like signal components, along with physiological regression to improve signal to noise ratio. Using these methods we found that CSF flow correlated with decreases in BOLD activity in 70 adults aged 19-76. We further found that brief, infrequent salient stimuli (visual oddballs, requiring a button press) were capable of provoking measurable CSF flow.

## Introduction

Flow of cerebrospinal fluid (CSF) is hypothesized to support waste clearance as a crucial aspect of the glymphatic system (1), the function of which is likely disrupted in Alzheimer’s disease (2). For this reason, measuring it has become an important goal of neuroscience. Changes in intracranial blood volume, which occur to dynamically support cognition, cause changes in CSF because the skull is rigid. Blood vessels dilate as neural activity increases, pushing CSF out of the foramen magnum, and constrict again when additional blood flow is no longer needed, drawing CSF back in. The bottom slice of fMRI acquisition, when it lies in a CSF reservoir, can be used to measure this process (3). CSF that has been outside of the volume of acquisition will have a different magnetic resonance signature from CSF further into the imaged volume, as a consequence of having been out of range of the radio frequency pulses. The measurement of this specific edge artifact allows identification of events that cause CSF inflow or outflow, and also to identify factors that may predict increased or decreased CSF flow (4).

Here we have analyzed CSF flow in 70 individuals across the adult lifespan who completed an oddball task with concurrent fMRI. This task was not designed to measure CSF flow, and the bottom acquisition slice does not cover the traditional fourth ventricle region used for most CSF analyses. Instead, it covers the spinal canal, within which CSF flow is visible in at least one prior report (5). Rather than rely on hand-tracing to identify the spinal canal, we propose an automated method for its identification. Furthermore, our data was acquired with a multi-echo EPI sequence, which allows the separation of BOLD-like signal components (which should not primarily represent CSF flow) from non-BOLD components, consistent with CSF flow. We further refine this high-confidence signal with physiological denoising. The oddball events in the task are very brief (∼70 ms), allowing a test of whether CSF flow is indeed evoked by short, salient events, and to test whether this flow differs across the adult lifespan.

## Methods

### Participants

Participants were enrolled in a study on healthy brain aging at Cornell University. They were screened for diagnosed cognitive impairment, neurological disease, severe head injury, ocular disease, cardiac arrhythmia, or any MRI contraindication, and had vision and hearing that were normal or correctable to normal. The study consisted of a visual oddball task with fMRI and other procedures reported elsewhere (6–10). The study was covered under Cornell Institutional Review Board protocol 1910009087 and participants gave written informed consent. Of 88 participants with complete fMRI data, our procedure to automatically identify CSF in the spinal canal (see below) failed in 5 (5.7%), leaving 83. Of the 83, 13 had no usable physiological recording, leaving 70 used in our final analysis (**Table 1**).

**Table 1.** Medians [interquartile range]. Kruskal-Wallis tests on whether each quantity differs between age groups.

| Group | n | Age | F/M | Spinal canal voxels, slice 1 (median, IQR) | Head motion (mean FD in mm, IQR) | Heart rate (mean bpm, IQR) | Respiration (breaths/min, IQR) |
|---|---|---|---|---|---|---|---|
| All | 70 | 19–76 | 56 / 14 | 58 [44–78] | 0.071 [0.046–0.132] | 69 [61–78] | 17.4 [14.4–20.3] |
| Younger | 28 | 19–44 | 21 / 7 | 58 [50–67] | 0.052 [0.037–0.098] | 70 [63–80] | 19.5 [16.5–20.8] |
| Middle | 24 | 46–63 | 19 / 5 | 62 [44–79] | 0.071 [0.047–0.113] | 68 [62–78] | 16.5 [13.3–18.6] |
| Older | 18 | 65–76 | 16 / 2 | 58 [41–83] | 0.115 [0.060–0.172] | 67 [57–77] | 16.1 [14.0–21.3] |
| Kruskal–Wallis p |  |  |  | .94 | .023 | .47 | .030 |

### Visual oddball task details

The oddball task had three types of stimuli presented for approximately 70 ms each (4 frames at 60 Hz): standard (60%, 1.8° visual angle purple circles isoluminant with the background), oddball (20%, 3.6° visual angle purple circles isoluminant with the background, requiring a button press), and bright (20%, standard-size purple circles brighter than the background). Between stimuli, participants fixated on a black dot (0.15° visual angle). The inter-stimulus interval was jittered (mean = 2.0 s, median = 1.5 s, range = 0.5-11.0 s, sequence designed with optseq2 (11)). Total task time: 10 min.

### MRI acquisition

We collected multi-echo echo planar imaging (EPI) using a GE Discovery MR750 3 T scanner with a 32-channel head coil at the Cornell MRI Facility (TR = 2.5 s; TEs = 14, 27, 40 ms; multiband factor 2; in-plane acceleration factor 3; 80° flip angle; 78 interleaved 2.0 mm slices; 21.5 cm field of view, acquired at a 72 × 72 matrix (≈3.0 mm in-plane) and reconstructed to 128 × 128 (1.68 × 1.68 × 2.0 mm voxels)). 240 volumes were acquired (600 s), and the first 2 volumes were discarded leaving 238 volumes (595 s) for analysis. Acquisition at this speed has previously been shown to allow CSF flow measurement (12). A T1-weighted MPRAGE sequence (TR = 2600 ms; TE = 2.85 ms; TI = 1000 ms; 7° flip angle; phase acceleration factor 2; 1.0 mm isotropic voxels, 200 slices) was also acquired.

### Automatic identification of spinal canal ROI

The spinal canal was identified automatically in each of the 4 lowest slices, on the raw echo 3 images, with the following procedure:

- A “start point”, meant to identify the single voxel at which the EPI signal was brightest over multiple slices, was identified. It was defined as the brightest voxel in the echo 3 images, across the whole run, averaged over the three lowest slices, within a central-posterior window of the field of view (i.e., excluding neck fat and vessels, which are brighter than CSF). This step was necessary to avoid selection of seed voxels that were noise (i.e., voxels bright in only one slice).
- In the lowest acquisition slice, the ROI was seeded at the brightest voxel within 10 mm of the designated “start point”. Up to 10 mm of deviation allows for correct identification of the spinal canal even if the acquisition slab is not fully perpendicular to it.
- In each subsequent slice, the ROI was seeded at the brightest voxel within 3 voxels of the ROI in the slice below.
- Within each slice, the ROI was then grown to the connected set of voxels, containing that slice’s seed, within 20 mm of the seed, whose intensity was at least 50% of the seed’s.

After this automatic process, we inspected each participant’s ROI slice by slice, blind to participant identity, to check visually whether the ROI lay on the spinal canal. Five participants failed QC at this step. In one participant, the 4th slice only was not usable; for that participant, slices 1-3 were used.

### Separation of S0 and R2* signals from EPI data

Within the spinal canal ROI, the mean signal S at each echo time TE (14, 27, and 40 ms) follows S = S0·e^(−TE·R2*)^, where S0 is the signal extrapolated to TE = 0 (not measured directly) and R2* is the rate at which the signal decays. To first order, small changes from volume to volume then follow ΔS/S = ΔS0/S0 − TE·ΔR2* (13,14). Changes in S0 alter the signal at every echo by the same fraction. Their sources include the inflow of fresh CSF, which has not recently been exposed to excitation pulses, into the lowest slices, as well as non-BOLD effects such as motion. Changes in R2*, by contrast, alter the fractional signal in proportion to TE. Their sources include BOLD-like effects, which arise from changes in the local concentration of deoxygenated hemoglobin. For each echo and each slice, the spinal canal ROI-mean signal was expressed at each time point as its fractional deviation from the run mean, with quadratic drift removed. Raw signals were used, without slice-timing correction, registration, or resampling. For each volume, ΔS0/S0 and ΔR2* were then estimated by fitting this model to the three echoes’ fractional deviations by ordinary least squares. The S0 component is reported as ΔS0/S0 in percent. The R2* component is reported as the percent signal change that ΔR2* would produce at the middle echo (-TE·ΔR2*, TE = 27 ms), so that positive values denote a BOLD-like signal increase. The same decomposition was applied to a tissue control region: FreeSurfer cerebellum and brainstem voxels (15) in the lowest slice containing at least 100 voxels of brainstem or cerebellar tissue, eroded by one voxel. Hereafter, the signals computed here are referred to as the S0 and R2* signals of each region (spinal canal or tissue).

### Physiological measurements

Heart rate and respiratory activity were recorded during fMRI at 100 Hz (photoplethysmography, PPG) and 25 Hz (respiratory, pneumatic belt) and used for denoising purposes. Systolic peaks were detected in the heart rate data with NeuroKit2 (16). Cardiac and respiratory phases for RETROICOR (17) were computed from the systolic peak times and from the pneumatic belt signal directly. Heart rate, from the inter-beat intervals, limited to plausible 30-180 bpm values, was convolved with the cardiac response function (18). Respiration volume per time was convolved with the respiratory response function (19). Recordings were used only if they covered the whole scan and passed visual inspection of a Poincaré plot of RR intervals.

### Nuisance regressors

The full nuisance regressor matrix for denoising included 5 Legendre polynomials to account for scanner drift, six motion parameters and their derivatives (12 parameters total), and 10 physiological regressors: eight RETROICOR regressors (17), plus heart rate·CRF and respiration volume·RRF. Additionally, outlier volumes were censored with AFNI 3dToutcount. The nuisance regressors were applied as follows (see **Fig. 1**): (A) To produce cleaned signals for spinal canal ROI S0 and R2*, tissue S0, and global BOLD. (B) Along with TENT regressors for oddball task events, described below, to identify responses to task events in those same tissues.

**Figure 1.**
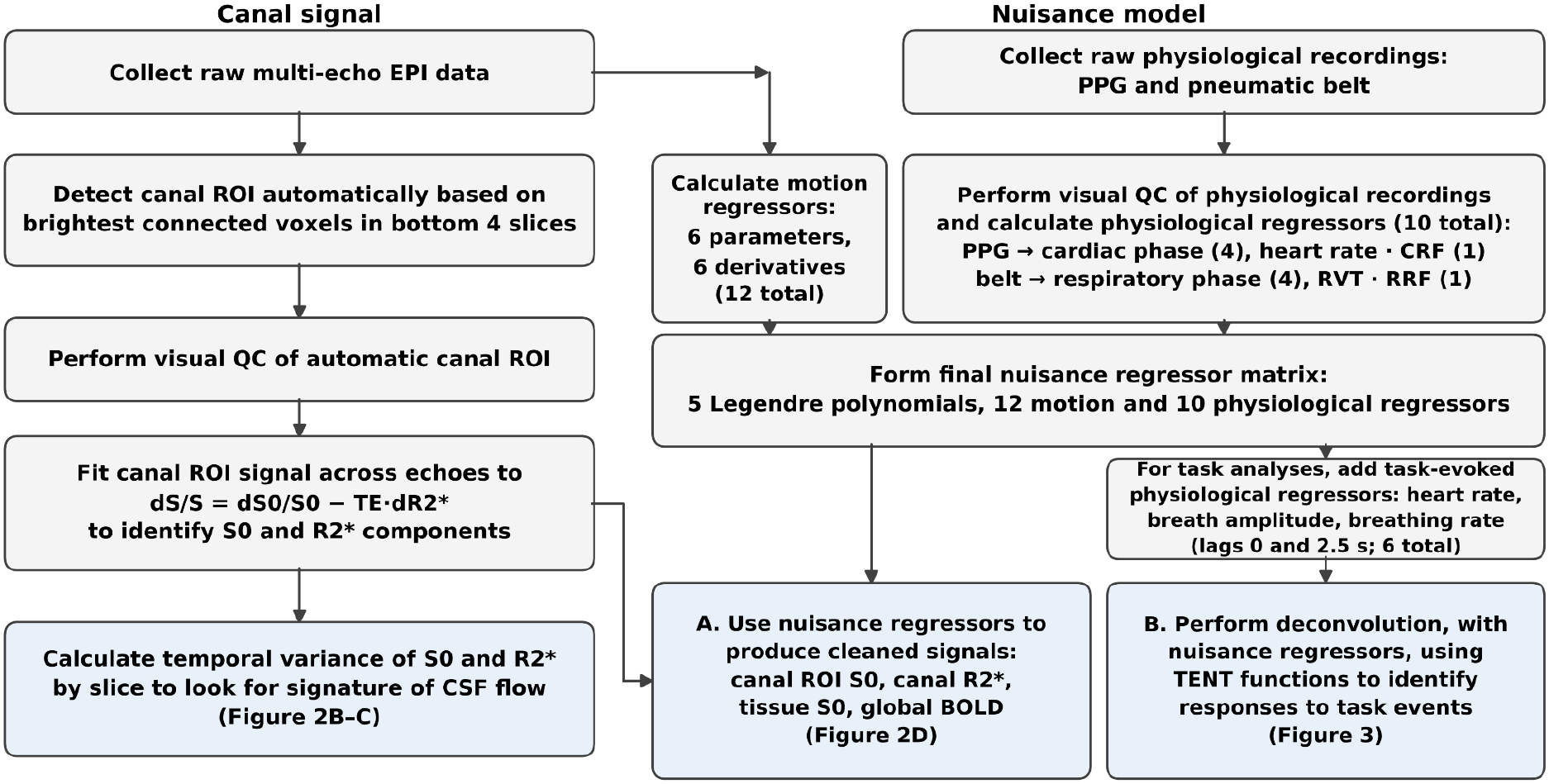
Flowchart of procedures. Procedures used to identify the spinal canal ROI, test for CSF flow, and test whether CSF flow is evoked by task events. ROI: region of interest; PPG: photoplethysmography; QC: quality control, EPI: echo planar imaging.

We found that together, the nuisance model explained a median 57% of spinal canal S0 variance (IQR 44-64%), with cardiac regressors explaining the most variance.

For all task-evoked analyses, we added 6 additional regressors to account for task-evoked physiological changes : raw heart rate, breath amplitude, and breath rate, each included twice, at a lag of 0 TR and 1 TR (20). All three were interpolated and sampled at the onset of each TR and z-scored within participant.

### Whole brain preprocessing and deconvolution

Whole-brain data were preprocessed with afni_proc.py (AFNI, (21)). Processing included removal of the first two volumes, despiking, slice-timing correction, alignment of the EPI to the T1-weighted image, volume registration to the minimum-outlier volume with resampling to 2.0 mm isotropic, brain masking, optimal combination of the three echoes and multi-echo ICA denoising with tedana (13,14), and scaling of each voxel to a mean of 100. The global BOLD signal was the mean of the denoised, scaled time series over all voxels in the brain mask. Correlations of spinal canal S0 signal with the negative temporal derivative (central difference) of the global BOLD time series were computed at lags of -1 to +2 TR, with positive lags denoting spinal canal S0 following the negative derivative of global BOLD (i.e., first BOLD changes, then spinal canal S0 changes).

Task-evoked responses were estimated by deconvolution. The slice 1 spinal canal S0 time series (and, as controls, spinal canal slice 1 R2*, tissue S0, and global BOLD) were regressed on the nuisance model with TENT basis functions (knots every 2.5 s from -2.5 to 15 s) for oddball, standard and bright stimuli. Oddball trials were split into within-participant quintiles by the size of their global BOLD response (mean cleaned global signal 5–7.5 s after onset minus the mean from −5 to 0 s).

To compare the CSF flow resulting from oddball-evoked BOLD responses, which included a button press, to CSF flow resulting from equally large BOLD responses that were not proximate to an oddball event, we identified, for each participant, the volumes with top 10% of cleaned global BOLD signal. We then divided them into events that occurred between 0-3 TRs (0 - 7.5 s) following an oddball event (N = 70, median 14 events per person), and those which occurred more than 5 TRs (12.5 s) after the last oddball event (N = 65; median 6 events per person, 5 participants had fewer than the three such events required for a per-participant average), and plotted the spinal canal S0 signal around each event. To test whether spinal canal S0 differed from zero around these events, we tested the mean spinal canal S0 in two windows against zero: during the rise of global BOLD (the TR preceding the event and the event TR; −2.5 and 0 s) and during its fall (the two TRs after the event; +2.5 and +5 s), with Holm correction over the two windows within each class. The two classes were compared by paired t tests, and framewise displacement around the same events was tested in the same way.

### Statistical tests

To test for differences between age groups (differences in physiological parameters, or in S0 variance across slices), Kruskal-Wallis tests were used to avoid assumptions of normality. To test whether the slice-1/slice-4 variance ratio was larger for S0 than for R2*, a paired Wilcoxon signed-rank test was used. To test whether correlations between spinal canal S0 signal (or any negative controls) and the negative derivative of global BOLD differed from zero, one-sample t tests on Fisher z transformed correlation values were used (for all participants, and separately within age groups). Spinal canal S0 and R2* coupling with the negative derivative of global BOLD were compared within participants by a paired t test on the Fisher z values.

To test whether task-evoked spinal canal S0 signals (or any negative controls) differed from zero at any time points, sign-flip max-T tests across the six knots from 0 to 12.5 s were used (5000 permutations). Within-age-group tests for significant effect at each knot were corrected for multiple comparisons at each lag (the different alignments of global BOLD and S0) by the same sign-flip max-T procedure. To test whether the magnitude of spinal canal S0 correlation with the negative temporal derivative of global BOLD, or the magnitude of spinal canal S0 responses to oddball trials, changed with age, the linear effect of age was tested by regressing each participant’s measure(s) on age. For the task-evoked responses, the age slopes at the six knots were corrected for multiple comparisons by permuting age (5000 permutations).

## Results

### Automatic identification of spinal canal ROIs

Spinal canal ROIs were successfully identified automatically (i.e., no manual tracing) in 94.3% of participants (83 of 88, see **Fig. 2A**); failures occurred in all three age groups (1 younger, 2 middle-aged, 2 older). The size of the resulting ROI did not differ by age group (**Table 1**). The median extent of the slice-1 ROI, from the most-anterior to most-posterior point, was 14.3 mm (IQR 11.8-15.1 mm). This was within the expected range for the spinal canal at C1 in healthy adults from normative MRI data (10.7–19.7 mm (22)).

**Figure 2.**
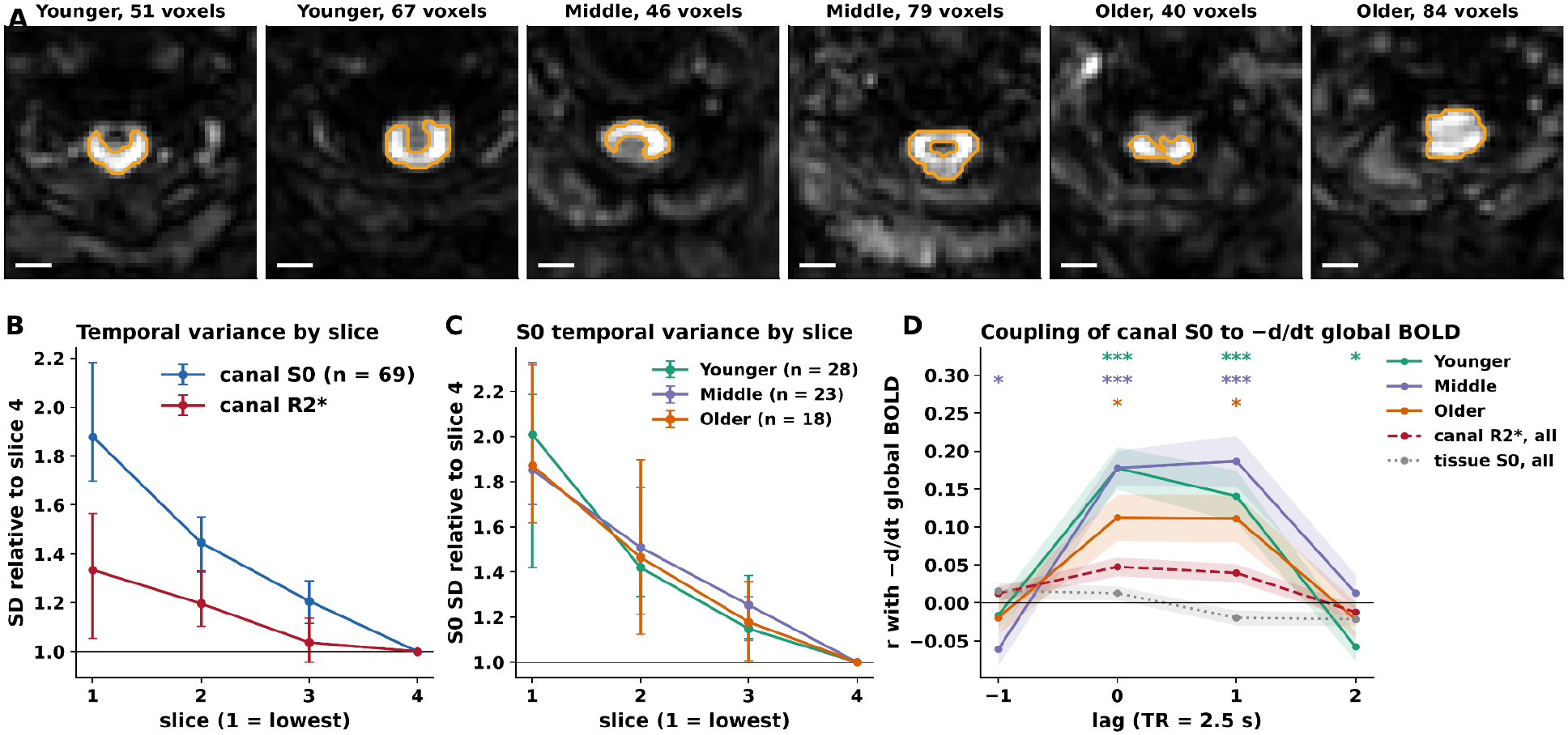
Signatures of CSF flow in the spinal canal. (A) Automatically identified spinal canal ROI (orange) in the lowest slice of two younger, two middle-aged and two older participants; scale bar 10 mm. (B) Temporal variability of the spinal canal S0 and R2* signals in the four lowest slices, relative to slice 4 (median and 95% CI, n = 69, 1 participant did not have slice 4 data). (C) The S0 temporal variability profile in each age group. (D) Correlation between spinal canal S0 and the negative temporal derivative of global BOLD at lags of -1 to +2 TR, by age group, with spinal canal R2* and tissue S0 as controls (mean ± SE). Stars: group mean differs from zero, corrected across lags (*p < .05, **p < .01, ***p < .001).

### Spinal canal CSF flow is visible in all age groups

The S0 signal within the spinal canal, the component of the EPI signal that is non-BOLD-like, was 1.88 times more variable in the bottom acquisition slice than in the fourth slice (**Fig. 2B**). Signal variability that systematically decreases as distance into the slab increases is consistent with CSF flow, because the greatest difference between CSF with unsaturated spins and CSF with saturated spins (the source of signal variability) occurs at the boundary of excitation. By contrast, the R2* signal, the component of the EPI signal that is BOLD-like, was only 1.33 times more variable in the bottom slice (S0 ratio was greater than R2* ratio in 72% of participants; paired Wilcoxon test on log ratios, p < .001, n = 69; one participant did not have slice 4 data). The S0 profile was similar in each age group (**Fig. 2C**; slice-1/slice-4 ratio Kruskal-Wallis p = .89).

Spinal canal S0 signal was significantly positively correlated with the negative temporal derivative of global BOLD at both 0 TR lag (r = +0.161, t(69) = 9.9, p < .001; **Fig. 2D**) and 1 TR lag (r = +0.149, t(69) = 7.5, p < .001; **Fig. 2D**). This is the expected signature of CSF flow (3). When global BOLD signal is decreasing, indicating reduced recruitment of blood flow, the decrease in cerebral blood volume draws CSF upward into the imaged volume. Within each age group separately, the coupling was significant at both lags (0 TR: younger r = +0.177, p < .001; middle-aged r = +0.178, p < .001; older r = +0.112, p = .013; 1 TR: p < .001, p < .001 and p = .015). The strength of coupling did not decline significantly with age (Fisher-z r regressed on age: -0.011 per decade, 95% CI -0.031 to +0.009, p = .29; spinal canal R2* control p = .90).

As negative controls, we tested whether the spinal canal R2* signal (BOLD-like signal) or tissue S0 signals (non-BOLD-like) were also correlated with the negative temporal derivative of global BOLD. Tissue S0 was not (r = +0.013, t(69) = 1.4, p = .16). Spinal canal R2* showed a coupling that was much weaker than that of spinal canal S0 (r = +0.048, t(69) = 3.8, p < .001; S0 versus R2*: paired t(69) = 6.3, p < .001).

### Spinal canal CSF outflow, and then inflow is induced by a brief visual oddball

We next tested whether global BOLD signals evoked by brief (∼70 ms) visual oddball stimuli were capable of provoking measurable CSF flow. Considering all participants together, we found that spinal canal S0 signal was significantly decreased (i.e., putative outflow, because no unsaturated spins entered from below the bottom acquisition slice, although outflow cannot be measured as directly as inflow; -0.57%, max-T p = .003) at 2.5 s after the oddball stimulus, and significantly increased (i.e., inflow, with unsaturated spins entering from below the bottom acquisition slice; +0.63%, max-T p = .010) at 7.5 s after the oddball stimulus (**Fig. 3A**). The global BOLD response peaked at 5 s and was lowest at 10 s, so the spinal canal signal extrema occurred 1 TR before the global BOLD extrema. Neither standard nor bright stimuli evoked a spinal canal S0 response that differed significantly from zero at any time point tested (smallest max-T p = .38 and .18). Similarly, neither spinal canal R2* nor tissue S0 signal differed significantly from zero at any timepoint tested (smallest max-T p = .40 and .85).

**Figure 3.**
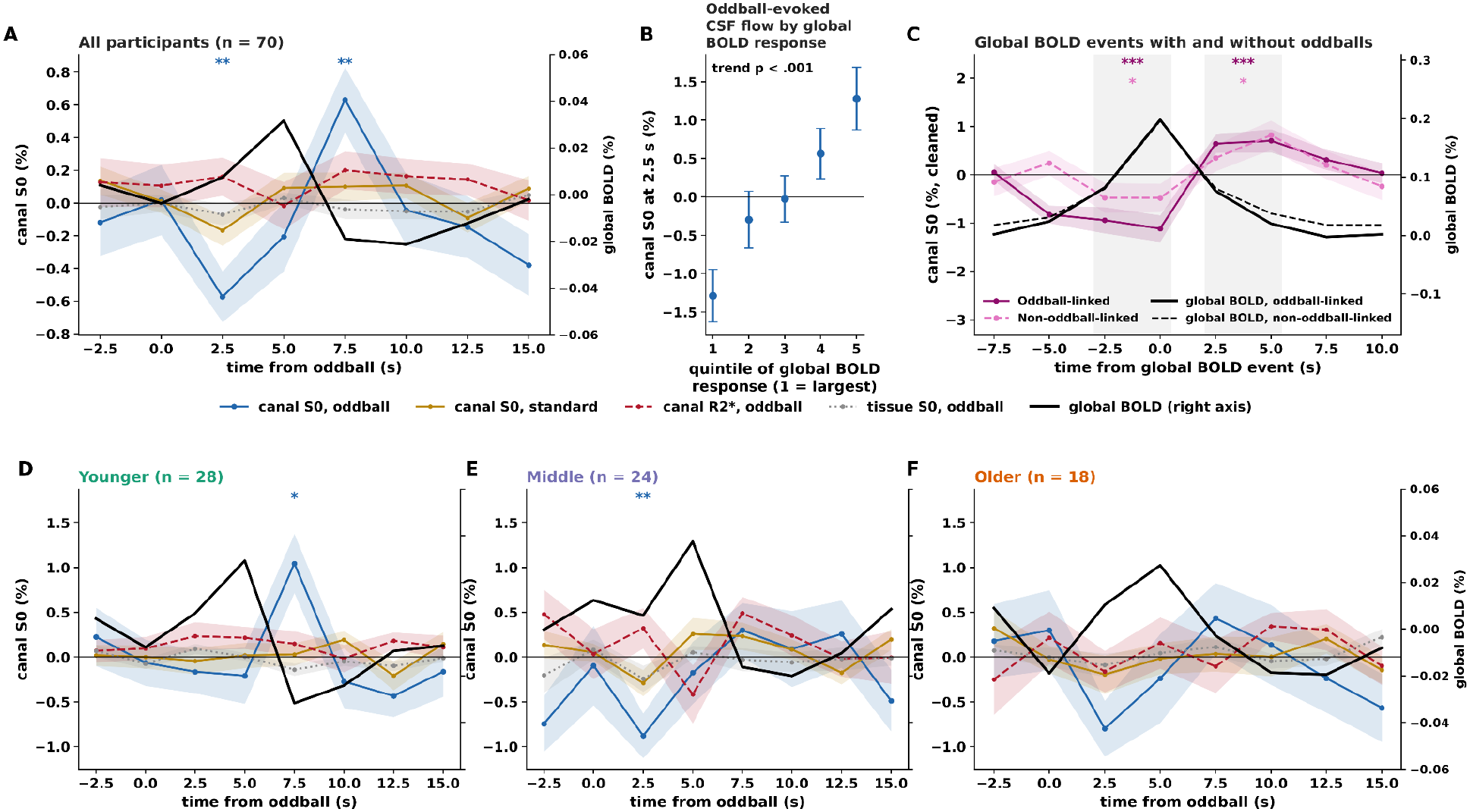
Task-evoked CSF flow. Deconvolved responses to oddball stimuli (mean ± SE across participants): spinal canal S0 (blue), spinal canal S0 to standard stimuli (gold), spinal canal R2* (red dashed), tissue S0 (grey dotted) and global BOLD (black, right axis), (A) in all participants and (D–F) in each age group. (B) Spinal canal S0 at 2.5 s after each oddball, relative to the pretrial baseline, by within-participant quintile of the trial’s global BOLD response (1 = largest). (C) Spinal canal S0 around global BOLD events that followed an oddball within 7.5 s (dark magenta, solid) or had no oddball in the preceding 12.5 s (pink, dashed), with the global BOLD signal for each class in black; shaded regions mark the two tested windows. Stars: spinal canal S0 differs from zero, corrected across time points (A, D–F) or across the two windows (C) (*p < .05, **p < .01, ***p < .001).

We next tested whether CSF flow evoked by oddballs was visible within each age group. Younger adults (N = 28) did not show a significant outflow signal at 2.5 s after the oddball stimulus (-0.16%, max-T p = .98), but they did show a significant inflow signal at 7.5 s (+1.05%, p = .020; **Fig. 3D**). Middle-aged adults (N = 24) showed only a significant outflow signal (-0.88% at 2.5 s, p = .004; **Fig. 3E**). Older adults (N = 18) showed neither at the corrected threshold (-0.80% at 2.5 s, p = .11; +0.43% at 7.5 s, p = .87; **Fig. 3F**). A linear effect of age on the spinal canal S0 response was not significant at any time point (all p > .25, corrected across time points).

To test for a dose-response relationship between global BOLD activity and CSF flow, we plotted the magnitude of evoked CSF outflow (the extent of which is not limited by skull volume), as S0 signal at 2.5 s after oddball events, by oddball response global BOLD quintile, and found a significant linear relationship between the size of global BOLD responses and the amount of CSF outflow at 2.5 s (+0.60 %/quintile, p < .001 **Fig. 3B**).

To test whether CSF flow followed global BOLD signal independently of button pressing, we plotted spinal canal S0 before and after two classes of global BOLD “events” (i.e., TRs with top 10% global BOLD signal). The first class of global BOLD events include those that occurred within 7.5 s of an oddball trial, which included button pressing. The second class included those that had occurred with no oddball in the preceding 12.5 s (**Fig. 3C**). For both classes, spinal canal S0 was significantly negative (i.e., outflow) averaged over the TR preceding the event and the event TR (oddball-linked: −1.03%, p < .001; unlinked: −0.48%, p = .042), and was significantly positive (i.e., inflow) averaged over the two TRs after the event (oddball-linked: +0.67%, p < .001; unlinked: +0.58%, p = .014, **Fig. 3C**). CSF flow did not differ significantly between classes (outflow: −0.53%, p = .087; inflow: +0.06%, p = .83). Framewise displacement at the time of the global BOLD event did not differ from each participant’s run average in either class (oddball-linked: −0.003 mm, p = .60; unlinked: +0.007 mm, p = .44), and the classes did not differ from each other (−0.008 mm, p = .37).

### Multi-echo decomposition is not required to detect the reported effects

We isolated the S0 signal component, and used spinal canal R2* and tissue S0 signals as controls, as a way to increase confidence in our measurements. However, many users do not have multi-echo acquisition. To determine whether isolating the S0 component was required to produce our main results, we repeated key analyses using a single echo (echo 2, TE = 27 ms). The slice-1/slice-4 variance ratio (2.05) and significant coupling with the negative temporal derivative of global BOLD (r = +0.18) still showed the signatures of CSF flow. Oddball-evoked outflow at 2.5 s was no longer significant, (−0.45%, max-T p = .16) but inflow at 7.5 s (+0.80%, p = .004) was. Multi-echo decomposition is therefore not required to detect most effects of CSF flow. It is required to attribute them to S0 rather than R2*, and to provide the controls we used.

## Discussion

We used a simple automatic process to identify the spinal canal from raw EPI data. This was successful in 94% of participants in an adult lifespan cohort. Within the automatic spinal canal ROI, we decomposed our multi-echo EPI signal into S0 and R2* components, and confirmed that the S0 component signal variability at the bottom edge of the acquisition volume was significantly greater than that of the R2* component. We then confirmed that the spinal canal S0 signal was significantly correlated, within all age groups, with the negative temporal derivative of global BOLD, while the tissue S0 signal was not. These two checks allowed us to be confident that we were observing CSF flow. We then measured whether CSF flow was evoked by oddball stimuli during our task, and found that within all participants, and within younger and middle-aged groups, it was. Although the coupling between global BOLD and CSF flow was similar across age groups, the global BOLD response and the resulting CSF inflow were numerically weaker in older adults and did not reach significance within that group, but could not be conclusively differentiated from younger adults.

The use of the spinal canal to observe CSF flow has recently been reported by others (5). Here, we add an automatic procedure to build usable spinal canal ROIs without tracing. This may facilitate this type of CSF flow analysis for researchers with fMRI volumes that incidentally cover the spinal canal, which is common when the full cerebellum is in the acquisition volume. For researchers who also have multi-echo acquisition, the techniques presented here should help to identify CSF flow simply and with high confidence by separating S0 and R2* signals, but we confirmed that the essential results are detectable without this step. The detection of CSF flow in response to task events in this analysis did, however, require careful physiological denoising, and was not detectable without it, since 57% of spinal canal S0 signal variance was explained by our physiological and motion regressors, particularly cardiac regressors, as previously reported (5).

The limitations of this work include the general limitation that these are indirect measurements of CSF flow, and particularly of outflow, which can only be measured as a lack of inflow. We were unable to measure CSF flow in the more traditional fourth ventricle due to the geometry of our scans. We lacked statistical power to test for age differences and also did not have, for comparison purposes, any stimuli which required a motor action (i.e., button press) without also being a salient target. Overall, this method does not allow us to differentiate between global BOLD changes that have autonomic vs. neural origins (20).

The stimuli used here were infrequent events, less than 100 ms, isoluminant, and covered only 3.6° visual angle, in contrast to the intense visual stimuli or color-dependent paradigms used in other reports of this phenomenon (23,24). Instead, they were distinguished by their identity as salient oddball trials requiring a button press. This work demonstrates that infrequent salient events are capable of producing CSF flow consistent with the size of the global BOLD response they provoke.

## Contributions

CRediT authorship contribution statement: **E. Riley:** Conceptualization, Methodology, Software, Validation, Formal analysis, Investigation, Resources, Data curation, Writing - original draft, Writing - review & editing, Visualization, Supervision, Project administration, Funding acquisition **H. Voss:** Validation, Writing - review & editing **E. De Rosa:** Resources, Supervision, Funding acquisition **A. Anderson:** Resources, Supervision, Funding acquisition, Writing - review & editing

## Data and code availability

All analysis code and de-identified data is publicly available at DOI 10.5281/zenodo.23989339

### Use of artificial intelligence tools

The initial data analysis code was written by the corresponding author between 2022 and 2026. Beginning in 2026, Anthropic models (Claude; models Opus 5.0/5.5, Fable 5.1) were utilized to optimize code, translate existing bash and R scripts into Python for faster performance and easier sharing, and implement and accurately report new statistical tests (permutation-based maximum statistic tests). To proofread the manuscript, the corresponding author used the same models to verify the accuracy of each number reported in the text against the analysis outputs and to populate data tables directly from model results to prevent transcription errors. AI was also used to organize all of the scripts that produce every number reported in this manuscript and place them within a shareable package. AI tools were not used to write the manuscript. The authors take full responsibility for the integrity and content of this work.

